# Assessing the Validity and Reliability of HD-DOT TD-fNIRS Resting-State Measurements in Rapid Succession Data Collection Settings

**DOI:** 10.1101/2024.03.04.583362

**Authors:** M. Parsa Oveisi, Davide Momi, Taha Morshedzadeh, Sorenza P. Bastiaens, John D. Griffiths

## Abstract

Functional magnetic resonance imaging (fMRI) has long been a cornerstone in the study of brain activity, but its high operational costs, limited availability, and restricted practical applicability have led researchers to seek alternative neuroimaging technologies. Recent advancements in high-density diffuse optical tomography (HD-DOT) and Time-Domain (TD) functional near-infrared spectroscopy (fNIRS) have emerged as promising solutions, offering the ability to generate detailed tomographic maps of hemodynamic fluctuations associated with neural activity. In this study, using the Kernel Flow device, we assess the performance of HD-DOT TD-fNIRS in terms of signal validity and reliability, particularly in rapid succession data collection settings. We conducted a multiple test-retest experiment involving fNIRS recordings from three participants across 20 ten-minute sessions of eyes-open resting-state brain activity over six days. Our findings indicate that HD-DOT TD-fNIRS systems like the Kernel Flow can reproduce hemodynamic patterns identified by fMRI, albeit with less spatial detail, and can detect resting state networks overall, though some individual network detections are not significant. The system performs consistently over days, with more variability within the time of day, and can capture subject-specific patterns with high accuracy as identified through FC fingerprinting analysis. We conclude that the new generation of HD-DOT TD-fNIRS systems holds significant promise for enhancing and expanding the measurement of functional brain data in both clinical and more naturalistic research settings. This study represents an important step towards a comprehensive understanding of the data quality and consistency achievable with these innovative neuroimaging devices.

## 1 Introduction

In recent years, functional near-infrared spectroscopy (fNIRS) has emerged as a promising neuroimaging technique, offering a non-invasive and portable approach to monitoring cerebral hemodynamics. By optically measuring changes in oxygenated and deoxygenated hemoglobin concentrations (Kim et al., 2017), fNIRS provides valuable insights into the neural correlates of cognitive and motor functions (Bauernfeind et al., 2016; Thomas and Nam, 2020). Its versatility and relatively low cost have facilitated the expansion of neuroimaging research beyond traditional laboratory settings, enabling studies in more ecologically valid environments. This flexibility aligns with our project’s aim to explore the potential of the Kernel Flow system, a wearable fNIRS device, in conducting neuroimaging studies outside the conventional fMRI laboratory setting.

fNIRS and fMRI, both critical in neuroimaging, have distinct features. fMRI offers high spatial resolution, crucial for detailed spatial brain mapping (Logothetis, 2008). Its limitation lies in the bulky and stationary setup, constraining use in naturalistic settings. In contrast, fNIRS is celebrated for its portability and flexibility, enabling studies in dynamic environments (Scholkmann et al., 2014). Additionally, fNRIS is capable of providing measurements in terms of absolute levels of blood oxygenation (Tak and Ye, 2014), which is an advantage over fMRI’s Blood Oxygen Level Dependent (BOLD) signal that primarily measures relative changes in blood oxygenation related to brain activity. This ability of fNIRS to measure absolute levels could potentially allow for a more direct assessment of cerebral oxygen metabolism and blood flow, without the need to infer these from relative changes alone. These differences render fNIRS and fMRI complementary, each fitting different research needs and environments.

HD-DOT (High-Density Diffuse Optical Tomography) in fNIRS is an advanced neuroimaging technique that employs high-density arrays of optical sensors to capture cortical hemodynamics with improved spatial resolution compared to conventional fNIRS. By using diffuse optical tomography algorithms, HD-DOT can reconstruct three-dimensional images of brain activity, enabling finer mapping of the cortical surface and the distinction of closely situated functional brain regions (Frijia et al., 2021). This higher density of optodes not only increases the penetration depth of near-infrared light, improving detection of deeper cortical activities, but also aids in more accurate three-dimensional reconstruction of brain activity. The sophisticated algorithms used in HD-DOT effectively reduce artifacts and enhance signal quality, making it particularly valuable for detailed brain activity studies and clinical research where precise localization is crucial (Uchitel et al., 2021). As such, HD-DOT stands out as a significant advancement in optical brain imaging, offering improved resolution and depth sensitivity compared to conventional fNIRS techniques.

Time-Domain functional Near-Infrared Spectroscopy (TD-fNIRS) is an enhanced form of fNIRS that employs ultra-short pulses of near-infrared light to measure brain activity (Torricelli et al., 2014). Its primary advantage lies in its ability to provide detailed information about the brain’s hemodynamic responses, including both the intensity and time-of-flight of photons. This feature allows TD-fNIRS to distinguish between signals from different tissue depths, greatly improving depth sensitivity and resolution (Ortega-Martinez, 2022). The time-resolved data also enhances the handling of artifacts, thereby improving signal quality (Wabnitz et al., 2020).

Recent advancements in fNIRS technology have led to the development of TD-fNIRS devices that integrate HD-DOT capabilities, offering enhanced spatial resolution and depth discrimination. One such example is the Kernel Flow system (Ban et al., 2022), which combines the advantages of TD-fNIRS and HD-DOT while addressing traditional limitations such as cost, complexity, and size. This compact and modular device provides dense channel coverage across the entire head, including the frontal, parietal, temporal, and occipital cortices, making it a significant step forward in making neuroimaging more accessible and practical for a wide range of applications.

The portability and ease of use of HD-DOT fNIRS devices like the Kernel Flow system enable neuroimaging studies to be conducted outside traditional fMRI laboratories, allowing for high-throughput data collection without extensive preparation. This flexibility opens up new possibilities for research designs that are more adaptable to naturalistic settings. Our study aims to explore the practicality and efficiency of using these devices in real-world environments, with a focus on maintaining data quality across continuous and successive use. We have implemented an intensive recording schedule to assess the performance of the Kernel Flow system under such conditions, examining the fidelity and consistency of functional connectivity measurements.

To assess the validity of HD-DOT fNIRS devices, we will evaluate their ability to replicate established hemodynamic spatial patterns previously identified in fMRI research. Specifically, we aim to determine whether devices like the Kernel Flow can accurately capture functional connectivity patterns, requiring high fidelity in recording brain activity and retaining precise spatial information. Our hypothesis is that the integration of TD-fNIRS and HD-DOT technology in these devices will enable them to successfully mirror these patterns, demonstrating their potential as reliable tools in neuroimaging research.

Assessing the reliability of HD-DOT fNIRS devices is also crucial. We will examine the consistency of their performance over various time scales, including hours and days, to ensure that they can produce stable and reproducible results under different conditions. This test-retest reliability assessment is essential for their application in longitudinal studies and repeated measures designs, highlighting the importance of reliability as a hallmark of a robust neuroimaging device.

## 2 Methods

### 2.1 Subjects

Three adult males (mean age = 32.66, SD = 4.49) who were part of our research team at Centre for Addiction and Mental Health (CAMH) were recruited for this study. Ethical approval for this study was obtained from the Research Ethics Board (REB) of CAMH. Informed consent was gathered prior to the experiments.

### 2.2 Experimental protocol

The protocol for this experiment, as summarized in Figure 1, was designed to evaluate the performance of HD-DOT TD-fNIRS devices in rapid succession data collection settings. The participants, three adult males, each underwent a total of 21 ten-minute eyes-open resting state recording sessions over a period of six days. The recording schedule was structured to include two days of dense multi-session recordings (6 and 7 sessions per participant, respectively) and four days of sparse two-session recordings per participant, with one session in the morning and one in the early evening. This intensive schedule allowed us to assess the reliability and consistency of the device across multiple sessions within a short time frame, providing insights into how these devices perform in settings that require rapid data collection.

**Figure 1:**
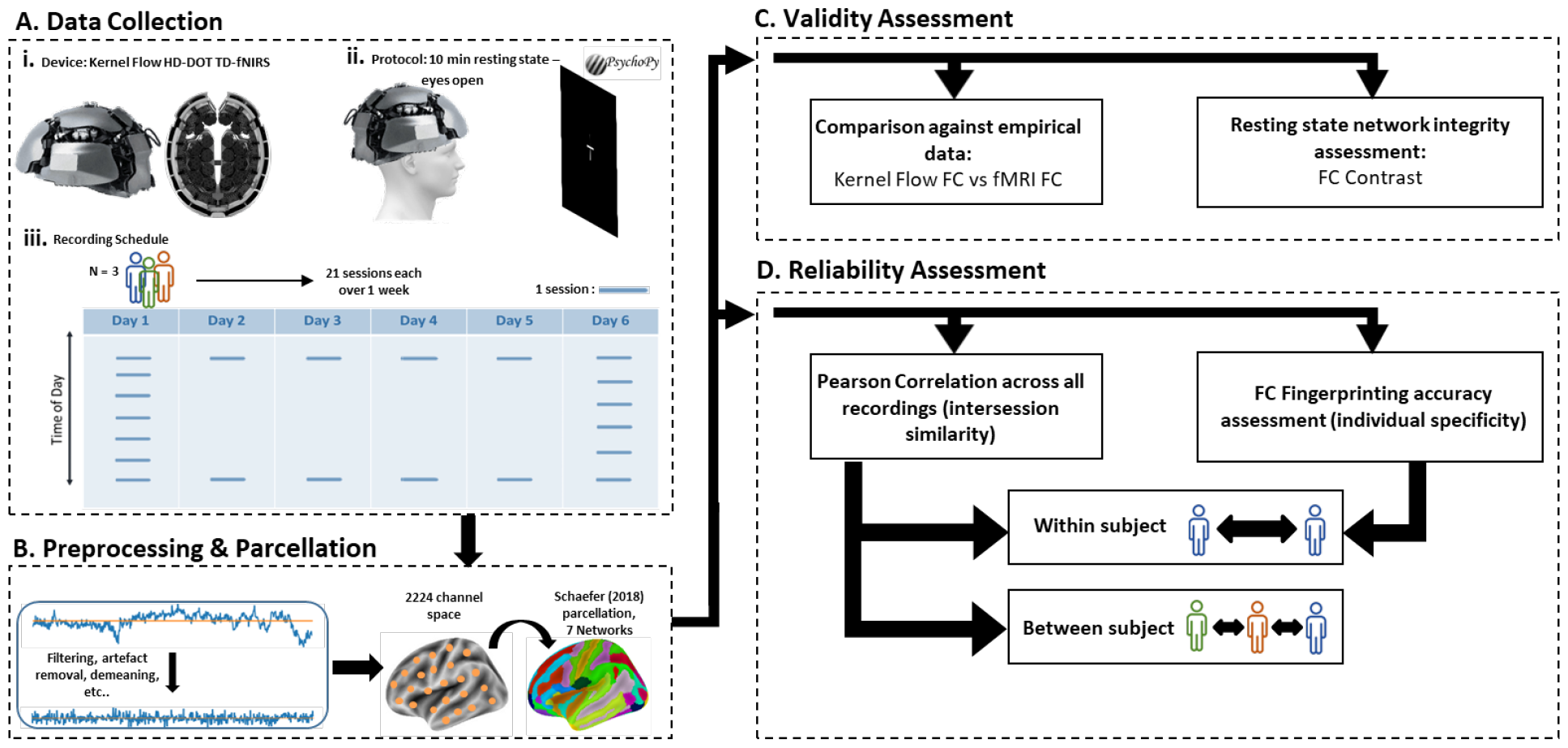
Summary of the experimental protocol and analyses. **A)** Eyes-open resting state fNIRS brain measurements recorded from participants (n=3) using the (i) Kernel Flow fNIRS device via (ii) 10-minute recording of participants staring at a fixation cross. (iii) Recordings were done over 6 days. Each participant underwent four days of light recordings (days 2-5) as well as two days of intense recording sessions (days 1 and 6) to assess time-of-day effects. **B)** The measurements were preprocessed, registered to brain space, and were parcellated based on Schaefer et al. (2018) brain atlas. **C)** Validity assessment of the Kernel Flow in identifying distributed brain activity by Functional connectivity (FC) matrix pattern analysis and Functional Connectivity Contrast analysis. **D)** Intersession consistency in capturing FC patterns was assessed via similarity comparison between FC matrices of all sessions and FC fingerprinting analysis.

During the recordings, the participants were instructed to remain seated and keep their gaze fixed on a white cross on a black LCD screen at an intermediate distance (approx. 70cm). The presentation of the fixation cross was programmed in the PsychoPy Python toolbox for psychology experiments (Peirce et al., 2019). The recordings took place in a quiet, dimly lit room. The participants were told to keep their eyes open during the recording, remain reasonably still, and not think about anything in particular. The wakefulness of the participants during the recordings were also monitored via a webcam placed below the LCD screen.

This experimental protocol was chosen to mimic real-world scenarios where neuroimaging devices may need to be used in rapid succession, such as in clinical settings or in studies involving multiple short-duration tasks. By assessing the validity and reliability of HD-DOT TD-fNIRS devices under these conditions, we aim to demonstrate their potential for broader application in neuroscience research and clinical practice.

### 2.3 fNIRS Recordings

The recordings were gathered via the Kernel Flow fNIRS device. For detailed specifications about the hardware please refer to Ban et al. (2022). In order to keep the configuration of the fNIRS device on the head consistent between subjects and between sessions, the device was mounted on the head so that it would be symmetrical and the frontal plate would rest 1cm above the eyebrows as an anchoring landmark.

### 2.4 Preprocessing

The data was collected and processed through the manufacturer’s internal pipeline which is comprised of the following steps: trimming edge artifacts, removing bad channels, calculating photon diffuse time-of-flight, conversion of light intensity metrics to absorption coefficients and hemoglobin concentrations, motion artifact correction using temporal derivative distribution repair (TDDR), and finally, short channel regression and converting optical intensity data recorded from fNIRS channels, to hemoglobin concentration data. (For more details, please refer to Ban et al. (2022).). This process resulted in a set of cleaned .snirf files for each recording. Additionally, the first 20 seconds of each recording were omitted to remove any potential interference due to movement and sound artifacts from initiation processes of the recordings and only the next 580 seconds were kept to account for sources of noise from termination of the recording and removal of the headset and to keep the length of recordings consistent between all sessions. A finite impulse response (FIR) filter-based lowpass filter was applied at 0.1Hz to remove low frequency fluctuations. Finally, the data was detrended using a 100s moving window average subtraction.

### 2.5 Functional Connectivity Analysis

Out of 63 recorded sessions, 5 recordings were identified as faulty and rejected and not included in further analyses. In order to examine distributed brain activity, the brain measurements (oxygenated hemoglobin measurements) were parcellated into 200 regions of interest (ROI) using the Schaefer et al. (2018) functional connectivity-based brain atlas, which also includes parcel membership to the 7 canonical Yeo et al. Yeo et al. (2011) resting state networks. However, the subcortical, insular, and other more basal regions were removed due to the fact that the fNIRS is not able to penetrate deep into the brain, leaving a total of 160 monitored brain parcels. The signals from each parcels were then z-score standardized such that the mean was shifted to zero and the signal was scaled to unit variance. The functional connectivity between parcels were assessed via calculation of the Pearson correlation coefficient between the time series of parcel signals using the SciPy toolbox in Python (Virtanen et al., 2020). All other handling of hemodynamic data was done using the Nilearn Python package (Abraham et al., 2014).

### 2.6 Functional connectivity contrast

We statistically compared the strength of functional connections and internal cohesion of parcels within each Schaefer Schaefer et al. (2018)network to parcels outside of the network. For this comparison we used the Functional Connectivity Contrast (FCC) metric (Kassinopoulos and Mitsis, 2022). FCC is a concept used in brain imaging studies to analyze large-scale networks. It involves assigning parcels, to specific networks and distinguishing between within-network edges (WNEs), which connect parcels within the same network, and between-network edges (BNEs), which connect parcels from different networks. The FCC quantifies the difference in correlation values between WNEs and BNEs. Higher FCC values suggest stronger positive correlations for WNEs compared to BNEs, indicating a more pronounced within-network connectivity. In our data, Fisher-Z transformations were applied to functional connectivity distributions (i.e., the Pearson correlation values from the functional connectivity matrices), and one-sided t-tests were conducted to evaluate the differences in internal connectivity strength of parcels within a given network, to their connection strengths to parcels outside of said network. This analysis is particularly relevant for our study as it provides a direct measure of the sensitivity of HD-DOT TD-fNIRS devices in capturing the integrity of resting-state networks. By employing FCC analysis, we aim to validate the ability of HD-DOT TD-fNIRS devices to accurately reflect the complex network dynamics of the brain, a crucial aspect for their application in neuroimaging research.

### 2.7 Reliability analysis

To assess reliability and consistency of the Kernel Flow in gathering measurements, similarities between the functional connectivity patterns of all recording were compared together. This was done by taking the lower triangle of the connectivity matrix for each session, unraveling the values into a one-dimensional array, and calculating the Pearson correlation coefficient of the array and the corresponding arrays for all of the sessions.

### 2.8 FC fingerprinting analysis

We employed FC fingerprinting (Finn et al., 2015) as a method to assess the reliability and individual specificity of the Kernel Flow fNIRS system. FC fingerprinting involves the comparison of functional connectivity matrices across different sessions for the same individual. The process entails identifying the most similar FC matrix for each recorded session and determining whether it corresponds to the same subject. This is achieved by two metrics: first, is the percentage accuracy of matching a session’s FC matrix with the average FC matrix of the corresponding subject. Second, is the percent prevalence that the most similar FC to a given session, also belongs to the same participant. The higher these values, the more reliable and individual-specific the FC patterns are deemed to be. Measures of similarity were the same as described in section 2.8.

### 2.9 fMRI dataset

To compare our fNIRS-based hemodynamic results to fMRI standards, we utilized a resting-state fMRI dataset with similar designs to our study. We used the Midnight Scan Club dataset from Gordon et al. (2017) (This data was obtained from the OpenNeuro database. Its accession number is ds000224). The fMRI data was acquired using a Siemens TRIO 3T MRI scanner. To align with our fNIRS data, we randomly selected data from three participants, each with 10 recordings taken one day apart. During the scans, subjects fixated on a white crosshair against a black background. Each session was truncated to 10 minutes for a direct comparison with our fNIRS recordings. The functional imaging employed a gradient-echo EPI sequence with specific parameters (TR = 2.2s, TE = 27ms, flip angle = 90, voxel size = 4mm x 4mm x 4mm). For additional details, please refer to the original study by Gordon et al. (2017).

## 3 Results

### 3.1 Validity results

#### 3.1.1 Functional connectivity patterns

As the change in oxygenated hemoglobin (HbO) levels are the most sensitive measurements to assess changes in cerebral blood flow (Scarpa et al., 2013), our analyses were based on HbO measurements. Figure 2A. shows the distributed activity patterns within the brain measured via Kernel Flow fNIRS and a reference eyes-open resting state fMRI dataset. The fNIRS-generated FC patterns show a number of qualitative similarities with fMRI-generated matrices. First, there is a strong pattern of high correlation along the main diagonal of both matrices. In this arrangement scheme of FC matrices, the high activity in the main diagonal line represents correlated activity of parcels within the same network, which are also mostly anatomically adjacent. Notably, the fMRI FC matrix shows a high activity pattern along the second diagonal as well, representing interhemispheric correlation of the same network parcels. In the fNIRS FC, however, this correlation is only restricted to the visual peripheral (VP) and dorsal attention (DA) networks (Fig. 2a, black arrow).

**Figure 2:**
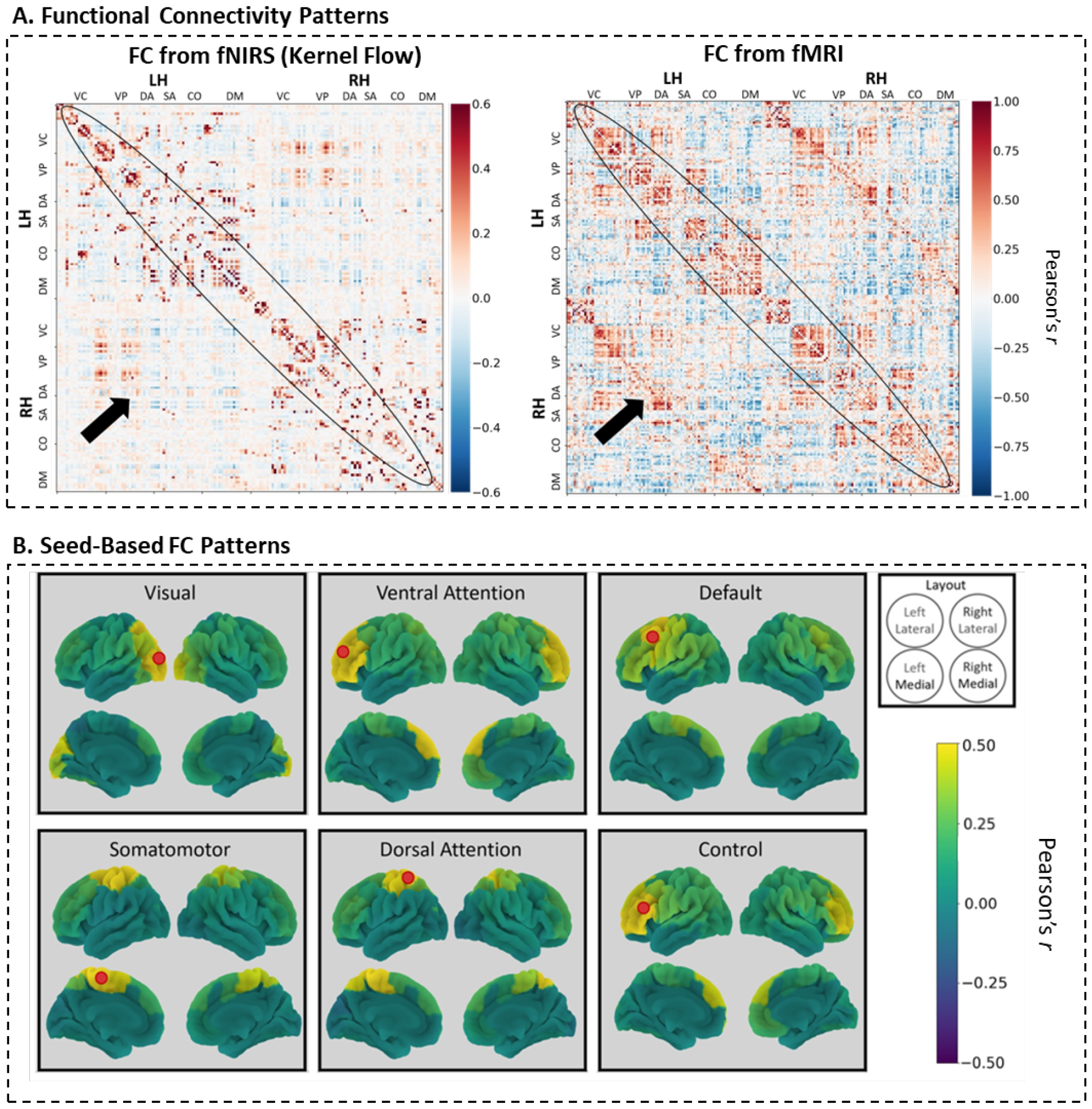
Ability of the Kernel Flow fNIRS device to identify canonical hemodynamic brain patterns. **A)** Functional connectivity matrices illustrating the correspondence between Kernel Flow fNIRS (left) and fMRI (right) data. The Kernel Flow matrix exhibits canonical features of functional connectivity, such as a strong diagonal activation (indicated by arrows) with both positive and negative correlations, and a trace presence of a secondary diagonal, indicative of cross-hemispheric connections. The fNIRS-derived matrix, while displaying these patterns, shows a lower resolution and reduced correlation strength compared to the fMRI-derived matrix, as evidenced by the bar scale values. **B)** Seed-based functional connectivity patterns from Kernel Flow fNIRS data for different brain networks, including Visual, Ventral Attention, Default, Somatomotor, Dorsal Attention, and Control networks. Each brain network exhibits contralateral activation, which is characteristic of hemodynamic responses in brain signal recordings. The lateral and medial views are provided for both left and right hemispheres, with connectivity strength indicated by the colour bar.

In regards to the details captured within the FC matrices, the parcellation resolution in the fNIRS FC matrices is lower, indicating a less detailed representation of brain activity. Additionally, the extent of correlations, as represented by the maximum correlation value (i.e., maximum value of the colour bar), is reduced in the fNIRS data. This suggests that while the Kernel Flow system can capture the overall structure of functional connectivity, the finer details and strength of these connections are less pronounced than in fMRI-derived patterns.

#### 3.1.2 Seed-based functional connectivity patterns

A key feature in hemodynamic data is the detection of cross-hemispheric correlations, which reflect the functional connectivity between analogous regions across the left and right hemispheres. To further validate the Kernel Flow’s effectiveness, we examined its capacity to identify these cross-hemispheric correlations (*r*) (Figure 2B.). We selected a reference seed in a representative parcel from each of the six detectable networks identified by Schaefer et al. (2018), excluding the limbic network due to its deeper cortical location and reduced accessibility to fNIRS light. Our analysis revealed significant correlations (r > 0.4) between parcels contralateral to the seed across all six networks, underscoring the Kernel Flow’s capability to capture essential cross-hemispheric functional connectivity.

#### 3.1.3 Resting state networks integrity

Figure 3 in our study presents the FCC analysis, which evaluates the internal correlations within each Schaefer et al. (2018) network against correlations with parcels outside of that network. Given that Schaefer networks are primarily defined as clusters of parcels with high internal correlations, our aim was to determine if the KF fNIRS system could detect these network-specific correlations. Our findings indicate that, on average, connections within each resting-state network exhibit higher internal correlations compared to between-network connections (Figure 3, first bar plot). However, this difference in correlation strength was not statistically significant for three out of the six resting-state networks examined. This suggests that while the KF system can identify internal network correlations, the distinction between within-network and between-network correlations may not be as pronounced as expected in some networks.

**Figure 3:**
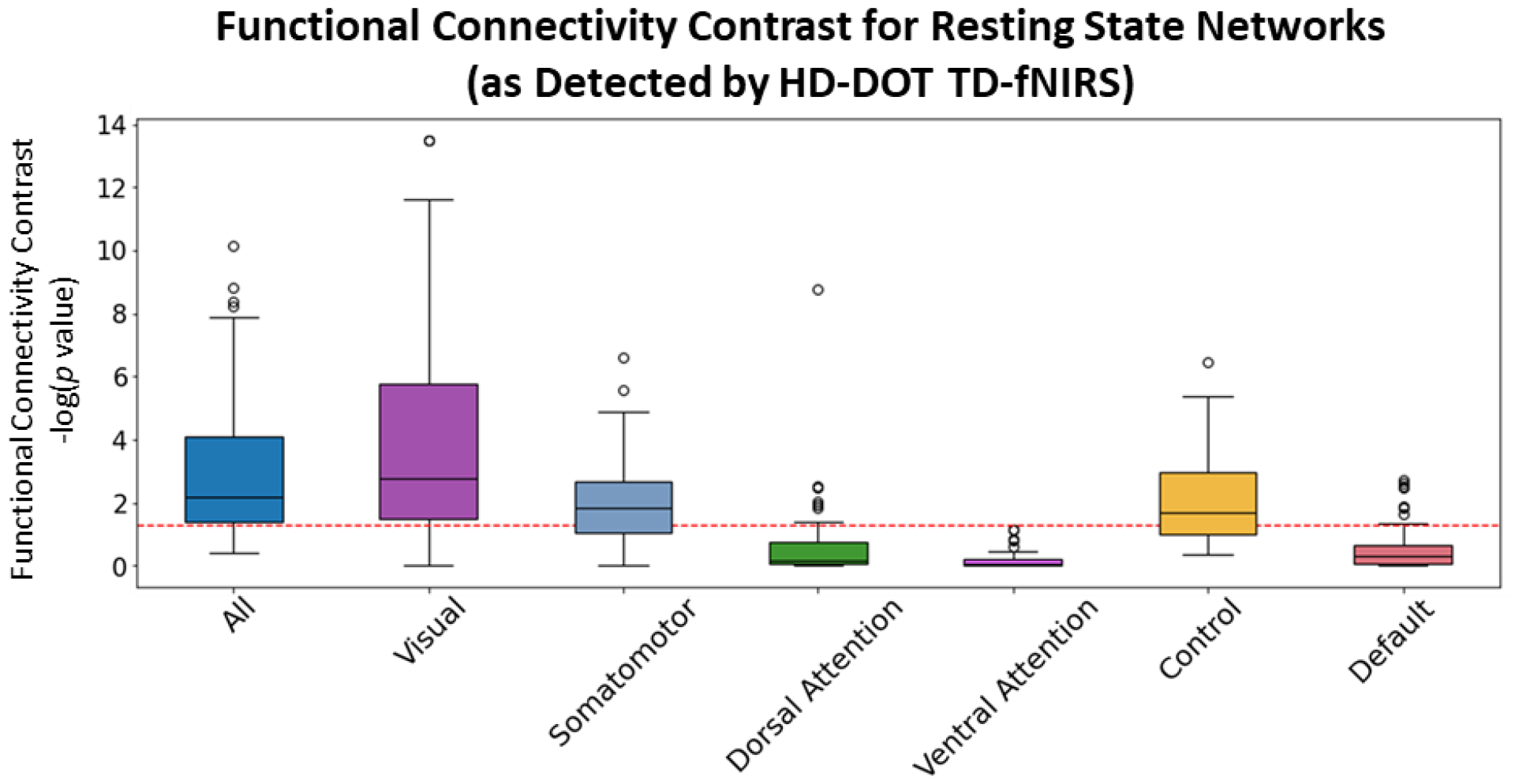
Within-network vs. between-network resting state network integrity as characterized by Functional Connectivity Contrast (FCC). The boxplot illustrates the Fisher z-transformed correlation values for functional connectivity within and between resting-state networks (higher is better). Higher functional connectivity is observed within individual networks compared to between different networks. However, this enhanced within-network connectivity is not statistically significant for half of the resting-state networks examined. The y-axis represents the negative logarithm of the p-value (-log10(p-value)), indicating the significance of the connectivity contrast, with values above the red dashed line (p < 0.05) considered statistically significant. Networks assessed include Visual, Somatomotor, Dorsal Attention, Ventral Attention, Control, and Default. Outliers are represented by circles. The lack of significance for three out of the six networks suggests variability in the degree of network segregation during resting-state conditions.

### 3.2 Reliability results

#### 3.2.1 FC pattern consistency

To examine the ability of the Kernel Flow in generating consistent measurements about distributed brain activity, we assessed the similarities between functional connectivity matrices of each session to every other. The results are depicted in Figure 4. A general observation is that all sessions are significantly correlated to each other (mean r>0.41, std=0.09). However, the experimental design utilized here allows us to examine the effects of factors that may influence the similarity between sessions, namely between-subject similarity vs. within-subject similarity over different recording sessions, and similarity over days and over different times of day. The results are summarized below.

**Figure 4:**
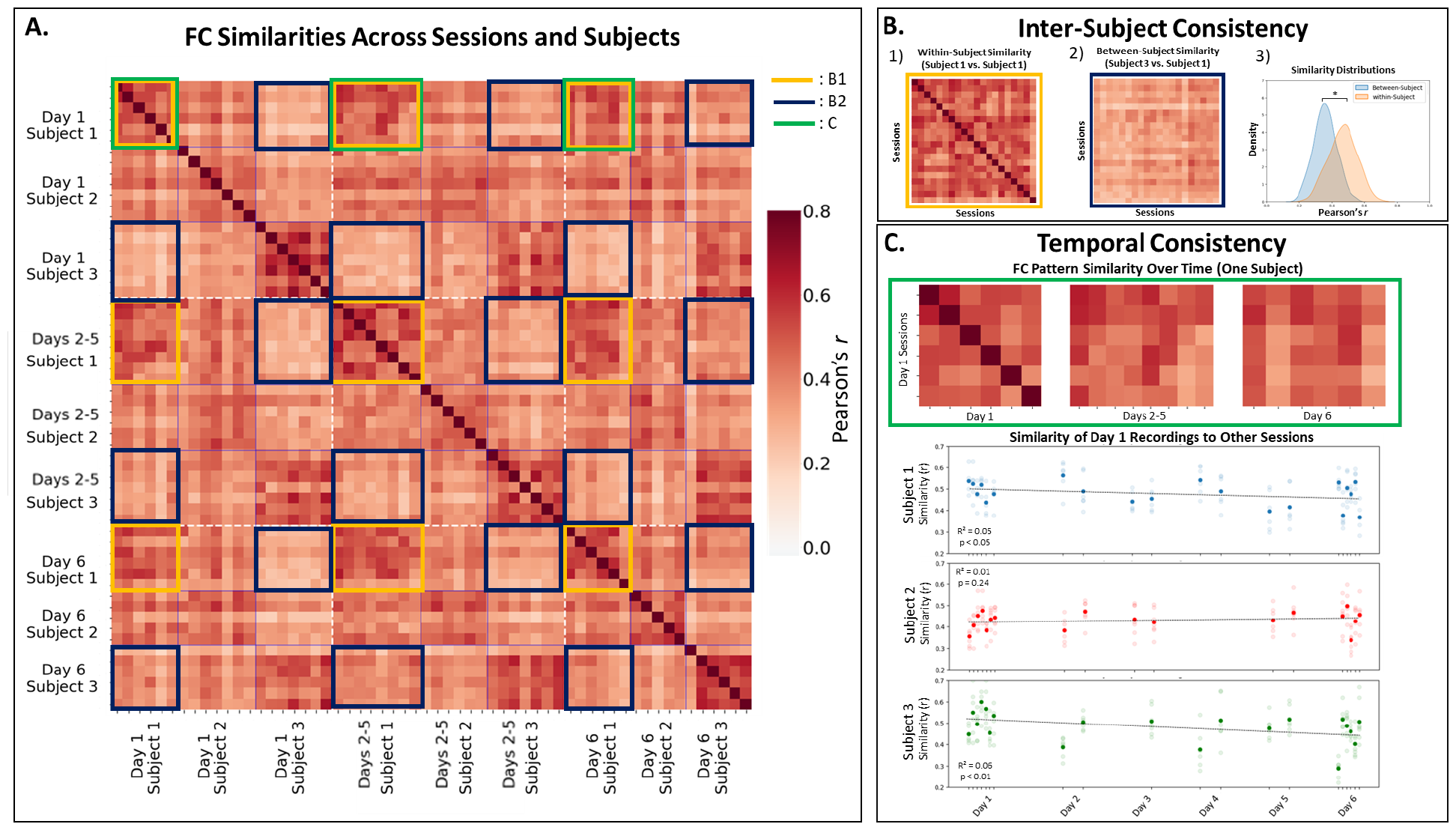
Temporal Stability and Inter-Subject Variability of Functional Connectivity Patterns. **A)**Comparison across all sessions and subjects. The heatmap is divided by white hashed lines indicating different days and blue solid lines delineating different subjects. Each tick mark represents a recording session at varying times of the day. The main diagonal blocks of high correlation values reflect strong within-subject FC pattern consistency during the same day. Secondary diagonals indicate moderate similarity of connectivity patterns for the same individual across different days. The off-diagonal blocks, showing lower correlation values, represent the least similarity and correspond to comparisons between different subjects at different times. This pattern highlights both the temporal stability of FC within individuals and the distinct FC signatures across different subjects. **B)** Inter-Subject Consistency. This segment of the figure illustrates the robustness of within-subject similarity in functional connectivity patterns compared to between-subject similarity. On the left, the heatmap shows a high degree of correlation in FC patterns when sessions from the same subject (Subject 1) are compared to each other, denoting strong within-subject consistency. In contrast, the middle heatmap, which compares sessions between two different subjects (Subject 3 vs. Subject 1), demonstrates notably lower correlation values, highlighting the individual uniqueness in FC patterns. The rightmost plot, a Kernel Density Estimation (KDE), visualizes the distribution of Pearson’s correlation coefficients within and between all subjects, statistically validating the consistency and significant distinctiveness of individual FC patterns. **A)** Temporal Consistency over hours and days. The top heatmap showcases FC pattern similarity over time for a single subject, with comparisons made between Day 1 sessions (dense recordings), Days 2-5 (sparse), and Day 6 (dense). The scatterplots below the heatmap represent the similarity of Day 1 recordings to other sessions for each of three subjects. Each plot reveals the variability and temporal trend of FC pattern similarity over the course of a week: Subject 1 exhibits a minor but significant decrease in FC similarity (R^2^=0.05, p<0.05), while Subject 2 maintains a stable FC pattern (R^2^=0.01, p=0.24), and Subject 3 shows a notable, significant decline (R^2^=0.06, p<0.01).Collectively, these visualizations provide insights into the temporal dynamics of FC patterns, revealing both the stability and the subtle changes that occur within individual subjects over a series of recordings.

#### 3.2.2 Reliability over subjects

High within-subject similarity is evident, as highlighted by the primary, secondary, and tertiary diagonal clusters in Figure 4A, indicating that individual functional connectivity profiles are more similar within subjects across sessions than between different subjects. The distributions of within-vs. between-subject similarity values demonstrate this distinction clearly (Fig 4B). Within-subject correlations had a mean *r* of 0.47 (std=0.09), whereas between-subject correlations were lower with a mean *r* of 0.36 (std=0.07). This difference is statistically significant (p<0.0001), underscoring the Kernel Flow’s capability to capture subject-specific functional connectivity patterns.

The results of our FC fingerprinting analysis also yielded high accuracies, indicating a robust capacity of the KF system to capture consistent and individual-specific FC patterns. Specifically, two key measures of accuracy were assessed: First, the success rate of matching a session’s FC matrix with the corresponding subject’s average FC matrix was found to be 96%. This high percentage demonstrates the system’s precision in identifying FC patterns unique to individual subjects across different sessions. A second analysis evaluated how often the most similar FC pattern for a subject corresponded to another session from the same subject. This measure also yielded a 96% accuracy rate. These results are significant as they indicate a high level of reliability in the Kernel Flow system’s ability to reproduce consistent functional connectivity patterns within subjects across different sessions.

#### 3.2.3 Reliability over time

The Kernel Flow’s performance was consistent across weekly sessions. However, a downward trend in similarity over time was noted, which was statistically significant for two subjects (Fig 4C.), indicating a minor decrease in similarity. Subject 2’s results did not show a significant change over time, implying stability in their FC patterns. The overall low R^2^ values suggest that the time factor’s impact is relatively small. Interestingly, high variability in similarity scores was particularly evident on days with dense recording schedules (days 1 and 6), as demonstrated by the larger spreads of similarity measurements within these days, indicating that the time of day may introduce more variability than the passage of days themselves.

## 4 Discussion

### 4.1 Overall Points

The Kernel Flow system demonstrated remarkable speed in setup and ease of use, requiring minimal training. We were able to record 63 recordings over a period of 6 days (and approx. 20 for high-intensity days), while only rejecting 5 sessions. Moreover, the recordings were done in a naturalistic setting with minimal spatial and range-of-motion restrictions. This aspect is crucial for its practical application in more diverse research or clinical settings, making it a convenient tool for both novice and experienced users.

### 4.2 Feasibility of Recording

The feasibility and high throughput capability of the Kernel Flow system represent a significant advancement in the field of neuroimaging. This system’s design allows for quick setup and ease of use, enabling researchers to conduct back-to-back sessions with minimal downtime. Such efficiency is unparalleled in traditional neuroimaging methods and opens new avenues for conducting studies with a high volume of data collection in short periods. This capability is particularly beneficial for studies requiring dense data collection schedules to capture dynamic brain activities or to monitor changes over time. By facilitating rapid and successive recordings, the Kernel Flow system offers a promising solution for large-scale studies, aiming to gather comprehensive datasets while maintaining participant comfort and engagement. This approach not only enhances the efficiency of data collection but also expands the potential for longitudinal and developmental studies, where frequent and reliable data capture is essential.

### 4.3 Validity

In terms of validity, the system showed good overall performance, particularly in terms of replicating canonical hemodynamic patterns. Although, in comparison to fMRI, the detection of these patterns were less robust. This disparity is expected, however, given the relatively lower spatial resolution of fNIRS, as well as limited penetration depth of fNIRS compared to fMRI (Scarapicchia et al., 2017). When analyzing functional connectivity matrices, we observed higher clustered activity within networks comprising larger parcels. Additionally, larger and more localized networks yielded better results, while networks with widespread distribution showed less significant outcomes. These variations could potentially be influenced by inconsistent helmet placement and the inherent differences in head shapes and sizes among participants. Notably, we utilized a fixed sensitivity matrix, which might have impacted these findings.

### 4.4 Reliability

Regarding reliability, the system exhibited robust performance. We found that individual similarities in brain activity were consistent across multiple sessions within the same day and across different days. Although the similarity decreased over longer time intervals, such as days, this was anticipated and aligns with the understanding that brain activities naturally vary over time. Importantly, the system effectively captured differences between individuals, showcasing high levels of unique pattern recognition and ’fingerprinting.’ This suggests its potential in identifying individual-specific brain activity. However, it remains unclear whether these unique patterns are predominantly due to the inherent brain signals of the individuals or influenced by factors such as head shape, signal intensity variations due to pigmentation, hair color, and hair shape.

### 4.5 Limitations and Future Directions

Several limitations were identified in our study. These include the variability in helmet placement and the homogeneity of our participant pool, which was limited in number and consisted solely of male subjects with pale skin. These factors could have influenced our results and limit the generalizability of our findings. Future research should aim to address these limitations, particularly focusing on accommodating variability in head shape and improving consistency in helmet positioning. Moreover, expanding the diversity of the participant pool will be crucial for validating the system’s applicability across a broader demographic spectrum.

## 5 Data and Resource Availability

The data supporting the findings of this study are available upon request from the corresponding authors. The code and analyses used in this study are available at the following GitHub repository: github.com/griffithslab/ Oveisi2024_kf-fc.

## 6 Acknowledgements

**Computing:** The computing infrastructure for this work included the CAMH Specialized Computing Cluster (SCC) and the cluster Narval at Calcul Québec, part of the Digital Research Alliance of Canada. The SCC is funded by the Canada Foundation for Innovation’s (CFI) Research Hospital Fund.

**Grants:** We gratefully acknowledge the financial support provided by the BranchOut Neurological Foundation for this research. The foundation’s funding was instrumental in facilitating the development and execution of our study.

## Author contributions *(in alphabetic order)*

**DM**: Methodology, Resources, Data Curation, Software, Resources. **JDG**: Conceptualization, Methodology, Software, Resources, Data Curation, Writing - Original Draft, Writing - Review & Editing, Visualization, Supervision, Project administration, Funding acquisition. **MPO**: Conceptualization, Methodology, Software, Validation, Formal analysis, Investigation, Resources, Data Acquisition, Data Curation, Writing - Original Draft, Writing - Review & Editing, Visualization. **SB**: Writing - Review & Editing, Visualization. **TM**: Writing - Review & Editing, Visualization.

